## Supplementary figures and images for "Pannexin 1 and Pannexin 3 differentially regulate the tumorigenic properties of cutaneous squamous cell carcinoma"

### Graphical Abstract

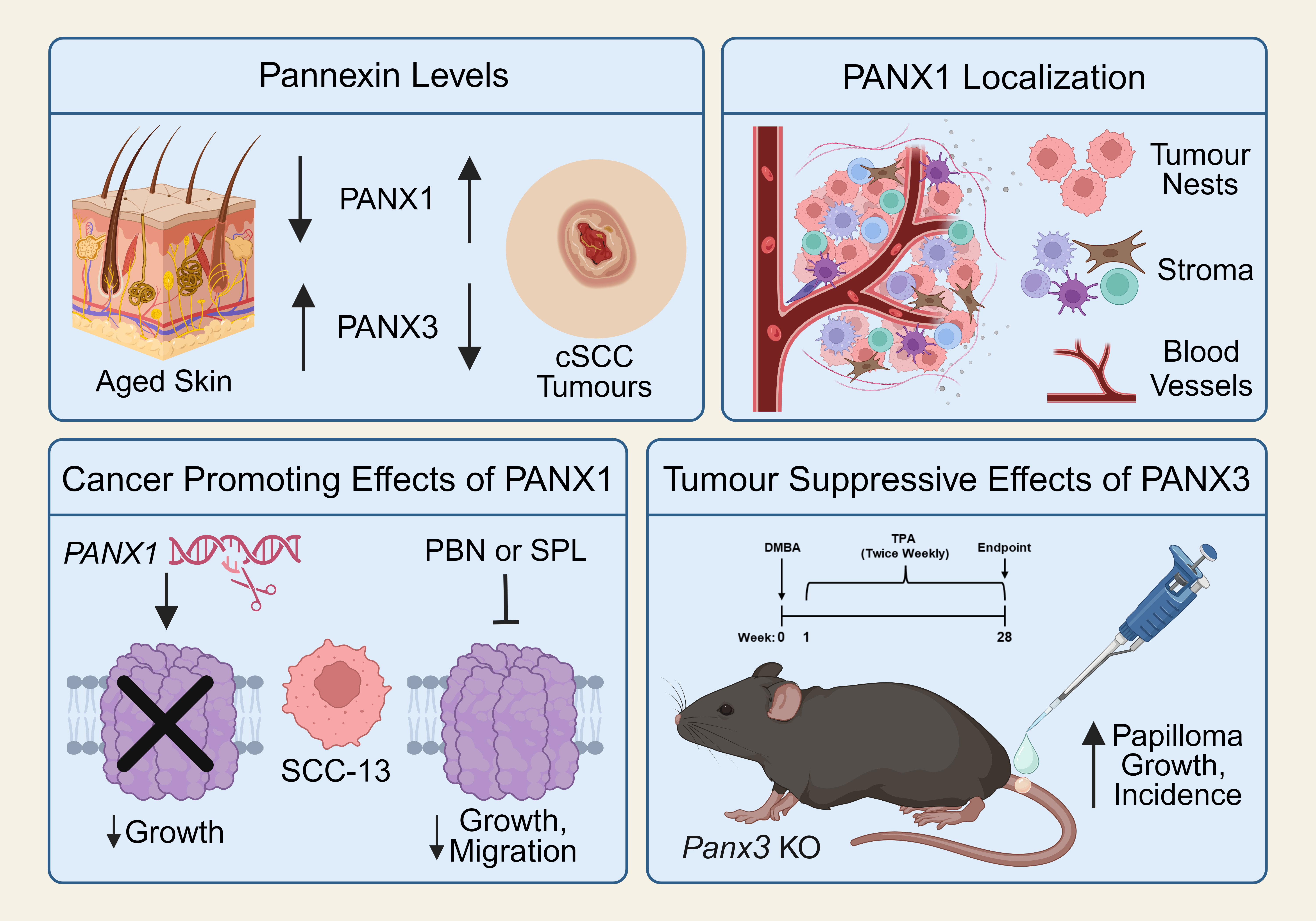
